# Dissociating the neural codes for multiple pitch perception in humans

**DOI:** 10.1101/2025.11.21.689749

**Authors:** Jackson E. Graves, Daniel R. Guest, Tess M. Starr, Angie Li, Anahita H. Mehta

## Abstract

Pitch is crucial for speech and music perception, yet its neural code remains contested. The classic ‘rate-place’ theory suggests that pitch is computed from spatial patterns of average neural firing rates along the cochlear tonotopic axis. These cues are thought to be robust for isolated harmonic complex tones (HCTs) but degraded for spectrally dense mixtures of simultaneous HCTs (e.g., musical chords) due to auditory filtering. In many cases, pitch perception remains feasible for HCT mixtures, but it is unclear whether listeners utilize residual rate-place cues or alternative neural codes. To adjudicate between these possibilities, we generated rate-place metamers (META), which are synthetic stimuli with simulated auditory-nerve average-rate responses that are nearly identical to those of original pitch-evoking stimuli (ORIG). Using these stimuli, listeners completed four psychophysical experiments that spanned a wide range of acoustic and cognitive complexity, from single-HCT pitch discrimination to harmonic-expectation judgments. Combining the behavioral data with predictive models of listener performance, we found that listener performance on single pitch discrimination tasks could be adequately explained by a rate-place model, with or without simultaneous pitch maskers (Experiments 1 and 2). More complex tasks involving attention to all pitches in a three-pitch mixture (Experiments 3 and 4) showed a pattern of results more difficult to account for with a rate-place model, perhaps suggesting a role for integration of temporal cues.

**Significance Statement:** Everyday listening typically involves parsing or integrating several simultaneous pitches, for example, while listening to music or conversations in noisy environments, rather than hearing a single, isolated pitch. Despite their ubiquity, the neural mechanisms for encoding multiple pitches are not well understood, which partly explains the limitations of modern assistive listening devices in noisy settings. To address this, we developed model-based synthetic stimuli to measure how distinct pitch cues contribute to pitch perception. This approach advances our understanding of the neural basis of music perception, with a goal of informing the design of more effective hearing prostheses.

## Introduction

Experimental approaches to understanding pitch perception date back to the 1800s (1). Many natural acoustic signals, such as animal vocalizations, human speech, and musical notes, comprise harmonic series of frequency components with a shared fundamental frequency (F0), typically evoking a perceived pitch corresponding to the F0. The limits of pitch perception for single harmonic complex tones (HCTs) have been well characterized behaviorally (2–5). However, in the natural world, listeners often encounter multiple, overlapping HCTs, especially in music, where simultaneous combinations of pitches are perceived without effort even by untrained listeners. Multiple simultaneous pitches also occur in group conversations, where each voice has a distinct pitch. Listeners can benefit from these voice-pitch cues to focus on talkers of interest (6) and receive suprasegmental information about talker emotion and intent (7). However, from the point of view of peripheral auditory physiology, it remains unclear what cues are utilized by the auditory system to parse these overlapping signals (see Figure 1 for examples of average auditory-nerve responses to a musical chord with three HCTs).

**Fig. 1.**
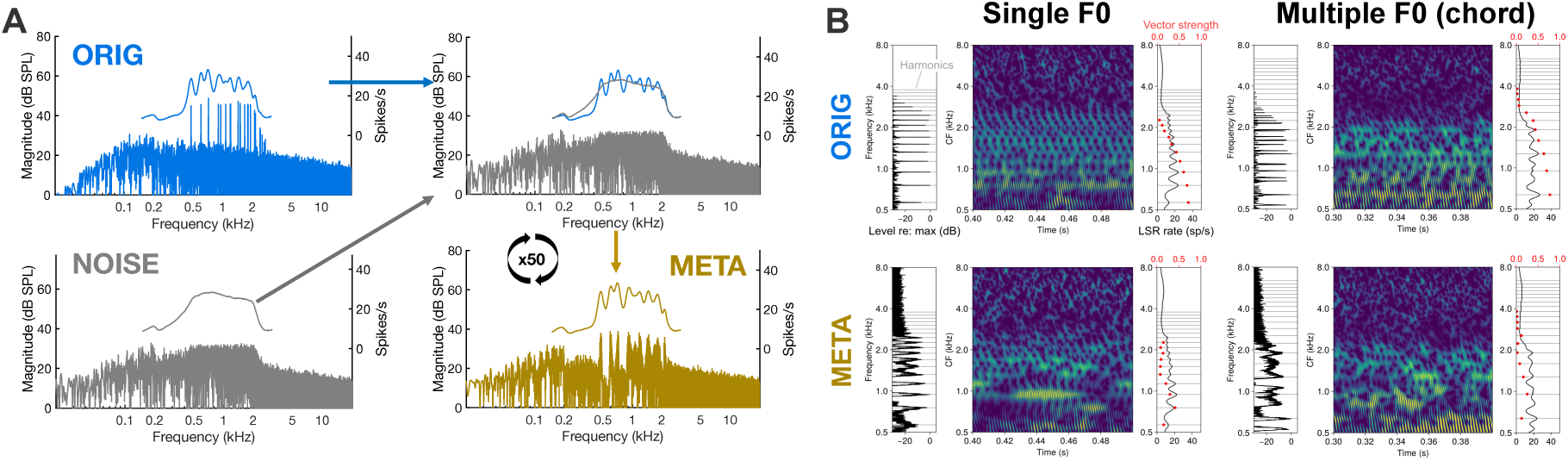
Stimulus generation. **(A)** Schematic representation of the processing pipeline for generating rate-place metamers. The magnitude spectrum of a Gaussian noise stimulus (NOISE) was iteratively modified so that its resulting auditory-nerve average-rate (ANAR) response, according to a model simulation, matched that of the target pitch-evoking stimulus (ORIG), resulting in a rate-place metamer (META). (**B)** Simulated auditory-nerve responses as neurograms, in response to single and multiple-F0 stimuli, in original and metamer conditions. Left: frequency spectra of each stimulus. Center: neurogram showing probability of firing by CF and time post stimulus onset. Right: rate-place profiles, shown as average firing rate at each CF. Red dots indicate vector strength.

Several possible neural codes that could support pitch perception have been identified based on human behavioral data and animal recordings (8–11). Classically, these include *rate-place codes*, where the pitch corresponding to the F0 is decoded from patterns of neural activity along the peripheral tonotopic axis elicited by the harmonic components in the HCT (Fig. 1B, right panels of subplots) (12–15), and *temporal codes*, where pitch is decoded from inter-spike intervals in phase-locked neural responses to temporal fine structure or the temporal envelope (Fig. 1B, red dots in right panels of subplots) (4, 16). More recently, interest has grown in codes that combine aspects of classical pitch codes, such as codes based on tonotopic spatial derivatives of phase-locked neural responses (17) or codes that employ place-specific use of phase-locked temporal cues (17–20). However, almost all this work has focused on single-pitch stimuli, such as isolated HCTs, while comparatively little is known about how listeners perceive pitch in more complex sounds, such as musical chords.

Adjudicating which neural codes are used by human listeners, and under what conditions, has proven elusive, particularly for the case of multiple-pitch stimuli (20, 21). Recently, computational modeling techniques have emerged as a key tool to address this question. For example, ideal observer analysis has been combined with computational auditory models to compare optimal performance based on auditory-nerve responses with actual human behavioral performance in frequency and F0 discrimination (20, 22, 23). While this approach can predict some trends in pitch behavior, such as the elevation of pure-tone frequency discrimination thresholds at high frequencies (20, 23), the technique is limited to a very small range of tasks for which an optimal solution can be derived analytically. More recently, researchers have investigated the conditions under which task-optimized deep neural networks (DNNs) demonstrate human-like pitch perception (24, 25). This approach is amenable to a wide variety of pitch stimuli and tasks, but it offers little clarity regarding which specific cues the model utilizes to achieve human-like performance.

Here, we propose an alternative approach using computational auditory models and novel synthetic stimuli we call *rate-place metamers* to dissociate the neural codes underlying pitch perception. To generate these stimuli, we iteratively modified samples of noise to produce synthetic stimuli with near-identical rate-place profiles (META) to original pitch-evoking stimuli (ORIG), based on an established model of the mammalian auditory nerve (26). Then, we compared behavioral measures of pitch perception for ORIG and META stimuli. As previously noted, rate-place theories of pitch posit that listeners use patterns of temporally averaged neural activity along the tonotopic axis — the *rate-place profile* — to perform pitch-perception tasks. If this is true, then we should expect that two stimuli that evoke the same rate-place profile (ORIG and META versions of stimuli) should also, to a first approximation, yield the same performance on pitch-perception tasks. Conversely, differences in performance between original pitch-evoking stimuli (ORIG) and their rate-place metamers (META) would suggest that an important element of the neural code is lost in metamerization, despite preserved rate-place cues, pointing towards importance of neural timing. Rate-place metamers are of particular interest for spectrally dense stimuli, such as mixtures of multiple simultaneous HCTs (i.e., chords). These stimuli are classically thought to have little useful rate-place information due to the limited resolving power of auditory filters (Fig. 1A) (21, 27). Some pitch tasks remain feasible under such conditions, but it is unclear whether listeners rely on weak residual rate-place cues or other available (presumably temporal) cues (20, 21).

In this study, we tested whether rate-place cues were sufficient for listeners to complete a series of tasks related to pitch processing in music (4, 16). Overall, we found that rate-place metamers supported pitch perception almost as well as original stimuli in simpler pitch tasks, such as discrimination of small changes in pitch, despite substantial degradation of temporal cues. In contrast, performance for more complex musical tasks, such as major-minor discrimination, differed for original and metamer stimuli, consistent with the notion that such tasks require integration of temporal cues Broadly, our results highlight the value of probing pitch perception across a wide range of stimulus and task complexity in elucidating the neural codes for pitch. Our results illustrate that assessing the contributions of the neural codes for pitch requires a nuanced understanding of the behavioral task that listeners are performing. Our experimental manipulations show that for the same complex stimulus, a single neural code might be sufficient for simpler pitch tasks and might require integration of more than one neural code for tasks that require a higher level of processing of multiple pitch stimuli, as commonly seen in musical tasks.

## Results

We conducted four behavioral experiments that ranged in stimulus complexity as well as ecological validity. The behavioral tasks were chosen to systematically tease apart the role of the rate-place code in the coding of complex pitch: pitch discrimination using a single HCT in isolation (Experiment 1), pitch discrimination of a single target HCT in the presence of two masking HCTs (Experiment 2), chord discrimination (Experiment 3) and subjective expectation ratings for chord sequences (Experiment 4). In order to measure behavioral limits of pitch perception separately from effects of F0 or spectral frequency alone, each task was conducted in different spectral regions and at different ranges of F0s (see Fig S1 for details and spectrograms of each frequency condition). All participants in each experiment completed tasks with both original (ORIG) pitch-evoking stimuli and their rate-place metamer (META) equivalents, in blocks presented in randomized order within each experiment.

The results of this series of experiments illustrate how different neural codes are weighted across tasks and frequency ranges. We also generated explicit model-based predictions for each task to evaluate the potential of rate-place and phase-locking information in explaining the behavioral data. Behavioral results from each experiment are shown in Figures 2-5, along with model predictions.

**Fig. 2.**
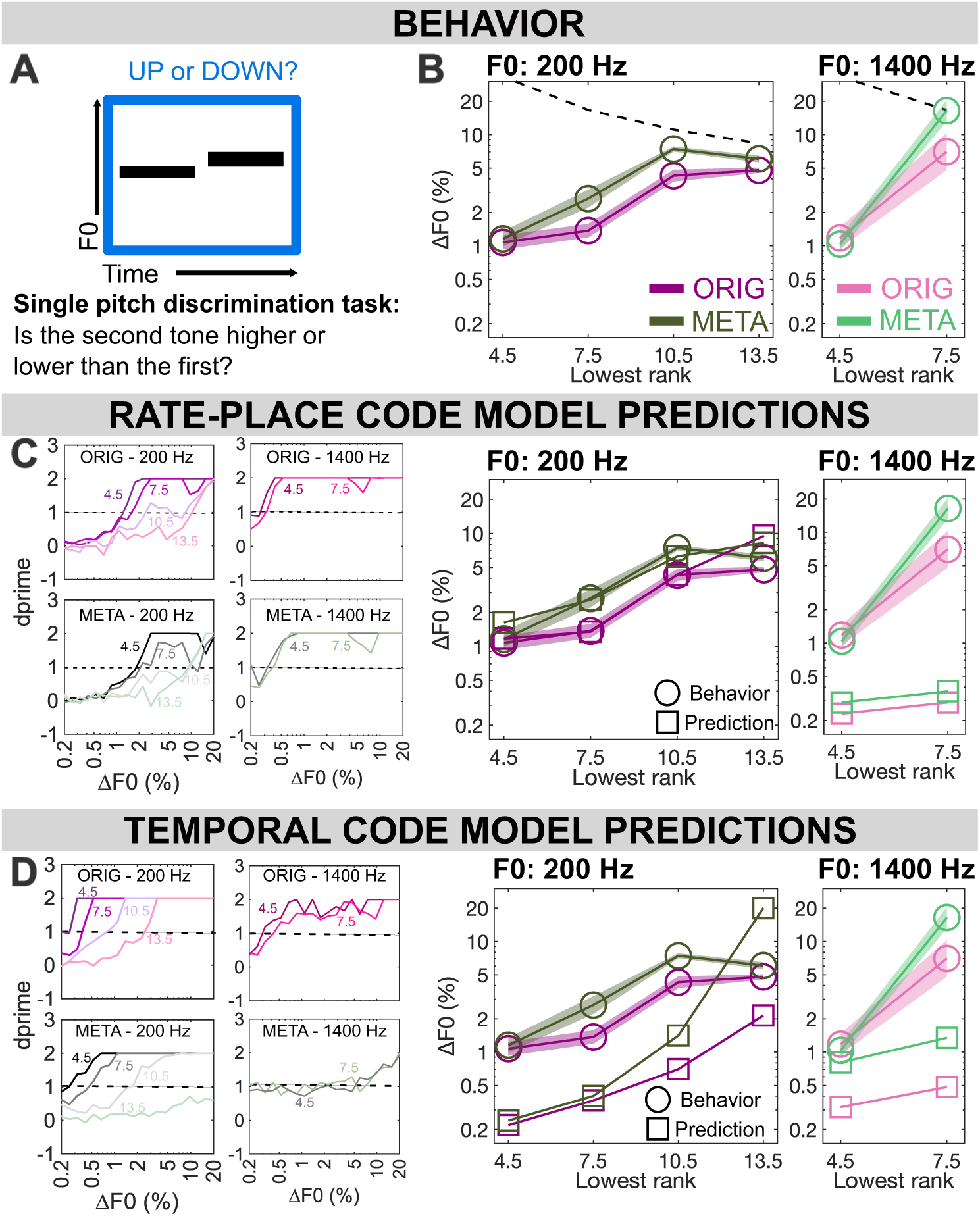
Results from Experiment 1: Single pitch discrimination. **(A)** Participants were asked to judge the direction of pitch change between two successive stimuli, either harmonic complex tones in noise (ORIG) or their rate-place metamers (META). **(B)** F0 difference limens by lowest harmonic rank, tone type, and F0 (N=21 for 200 Hz, N=11 for 1400 Hz, due to high-frequency hearing screening). **(C)** Predictions from a rate-place model, with neurometric functions by condition on the left and corresponding thresholds on the right, plotted against human behavioral thresholds. RMSe shows the root-mean-square error between model and behavior by condition. **(D)** Predictions from a temporal model, as in (C) but using summary autocorrelation functions rather than rate-place profiles. Throughout: shaded regions ± 1 SEM.

### Experiment 1: Single-pitch discrimination

In our first test of rate-place model-based stimuli, we compared listeners’ performance on F0 discrimination of individual HCTs, a task that has previously been used (28) to investigate how pitch discrimination thresholds change as a function of resolvability and fundamental frequency. As indicated in Figure 1B, harmonic components of lower-rank (harmonic rank <10) yield distinct rate place cues in the periphery. HCTs including lower numbered harmonics have better pitch discrimination thresholds than HCTs consisting only of higher numbered harmonics (harmonic rank >10), where the pitch perception is based solely on temporal envelope cues.

### Pitch of rate-place metamers is less discriminable in some conditions

In Experiment 1 (n = 21 for f0=200 Hz; n=11 for f0=1400 Hz), results for ORIG replicate the findings of Mehta and Oxenham (2020): for F0s of 200 Hz, F0DLs transition from good performance (near 1% ΔF0) to poor performance (near 5% ΔF0) as the lowest included harmonic rank shifts from the 7th-8th to the 10th-11th. A repeated measures ANOVA was performed to compare the effect of stimulus type (original vs. metamers) and lowest harmonic rank (4.5, 7.5, 10.5, 13.5 kHz) on participant performance in Experiment 1. Results indicated a significant main effect of stimulus type (F(_1,20_)=60.733, p<0.001), a significant main effect of condition (F(_3,60_)=109.706, p<.001) as well as an interaction (F(_3, 20_)=9.886, p<0.001). Post-hoc comparisons with Bonferroni correction indicate no significant difference of stimulus type for harmonic rank 4.5 (p=1.000) or harmonic rank 13.5 (p=0.174), but significantly poorer thresholds for META at harmonic rank 7.5 (p<0.001) and 10.5 (p=0.008). Thresholds have been thought to be solely based on rate place cues for harmonics below rank 10, however, the elevated thresholds for META at 7.5 suggest that listeners might be using more than only rate place cues for this frequency region. Using this metamerization approach to isolate neural cues allows us to get a better understanding of these phenomena, thought to be well established in pitch perception.

However, for the higher F0 of 1400 Hz, this same transition in behavioral performance happens earlier, as lowest rank shifts from 4th-5th to 7th-8th (consistent with prior evidence (28)). These transitions happened at the same point for META and ORIG. However, thresholds for META are elevated relative to ORIG thresholds for unresolved (lowest rank ≥10) pitch. A repeated measures ANOVA was performed to compare the effect of stimulus type (original vs. metamers) and lowest harmonic rank (4.5 and 7.5) on the high-frequency (1400 Hz) results from Experiment 1. Results indicated no significant main effect of stimulus type, (F(_1,10_)=1.716, p=0.220), but an effect was shown for lowest harmonic cutoff (F(_1,10_)=66.444, p<.001) as well as an interaction (F(_1,10_)=5.071,p<0.048). Post-hoc comparisons (Bonferroni corrected) indicated no effect of stimulus type at harmonic ranks 4.5 (p=1.000) or 7.5 (p=0.481). Comparison of root-mean-square error (RMSe) of model predictions against behavioral results revealed that the rate-place model predictions more closely matched behavior results (average RMSe across conditions = 1.049) compared to the temporal model (average RMSe across conditions = 2.424). See supplemental methods (Table 1) for condition-by-condition RMSe results.

### Single-pitch discrimination suggests dominance of rate-place cues

Based on model predictions seen in Figure 2C, threshold predictions for both ORIG and META based on the rate-place code provides a reasonable match to the data for F0=200 Hz, though it is notable that this model underestimates human performance for unresolved pitch (when lowest rank is 13.5). This is to be expected as humans are thought to rely on temporal cues in such conditions, whereas the rate-place model predictions were constrained to only use rate-place cues. It is also notable that the differences in behavior between ORIG and META are predicted in the threshold estimates using a rate-place model. This is likely due to variability in the rate-place cues provided by individual stimulus-length (1s) samples of metamers, which were matched to ORIG stimuli at the level of the auditory nerve on a 10-s timescale but may vary from second to second (see metamer construction details in Methods).

On the other hand, thresholds predicted from the outputs of a temporal model based on autocorrelation (Figure 2D), show overestimation of performance for F0 = 200 Hz, a typical result for this class of model. It also predicts a large gap between ORIG and META for unresolved low-F0 performance which is not observed in behavior, with this gap only emerging at unresolved regions (lowest ranks 10.5 and 13.5). This result is consistent with the qualitative difference in the strength of phase locking to temporal fine structure in the ORIG and META stimuli (Figure 1B).

Both models overestimate human performance for high F0s in the 1400 Hz range. For the rate-place model, the difference between low- and high-F0 thresholds predicted here (∼4x better for high) agrees with a previous ideal-observer modeling study (26) and reflects the (relatively) sharper peripheral filters at high CFs in the peripheral model. However, this discrepancy may not ultimately be relevant, since high-frequency pitch discrimination may be largely constrained by non-peripheral mechanisms (29, 30). The rate-place model predicts a small gap between ORIG and META at these frequencies, while the temporal model predicts a large gap. Behavioral results show no gap at 4.5 and a moderate gap at 7.5. Overall neither model fully captures the pattern of human results. This discrepancy could also stem from the auditory model not being optimized at these higher frequency ranges for humans.

### Experiment 2: Hearing out a single pitch in a mixture

We next sought to determine whether listeners could achieve similar pitch discrimination performance for ORIG and META stimuli even when presented with concurrent maskers. It has been previously shown that listeners are able to parse out a single pitch from a mixture of three HCTs behaviorally (21). For the single-pitch F0DLs in Experiment 1, we found that the rate-place code was sufficient to explain the behavioral results. However, because concurrent masker HCTs degrade target rate-place cues (Figure 1B and Figure S1), it is often thought that a rate-place code cannot generalize to more acoustically complex multiple-pitch stimuli. To investigate this further, we measured F0DLs for a single pitch within a triad (i.e., a combination of three simultaneous F0s).

### Rate-place metamers are largely equivalent to original stimuli when hearing out a single pitch in a mixture

Overall, F0DLs in Experiment 2 are higher compared to Experiment 1 for both ORIG and META conditions, consistent with previous findings (21). Despite masking effects from concurrent masker HCTs, F0DLs for the target HCT follow the same pattern as a function of cutoff frequency for META and ORIG (Figure 3B). We performed repeated measures ANOVA for each frequency range separately to compare the effect of stimulus type (ORIG vs. META) and lowest frequency cutoff for each of the 3 F0 conditions. Supplementary figure S1 shows the average rate place information for the various frequency cutoffs and F0s. For the low frequency conditions (F0 = 112.5 Hz), no significant main effect of stimulus type was found (F(_1,20_)=1.970, p=0.176) but results did indicate a significant main effect of lowest cut off (F(_3,60_)=12.345, p<.001) as well an interaction (F(_3,60_)=2.962, p=0.039). Post-hoc comparisons showed no difference of stimulus type at any frequency cut off (all cut-off condition comparisons were p=1.00 except 1.5 kHz (p=0.478)). For the mid-frequency conditions (F0 = 300 Hz), we found a significant main effect of stimulus type (F(_1,20_)=17.078, p<0.001), as well as lowest cut off (F(_6,120_)=47.900, p<0.001) but not a significant interaction (F(_6,120_)=2.065, p=0.062). Post-hoc comparisons (Bonferroni corrected) found no effect of stimulus type at any cut-off conditions (0.5, 0.7, 1.5, 3, 4 kHz (p=1.00), 1 kHz (p=0.603) or 2 kHz (p=0.341)). For the high-frequency conditions (F0 = 900 Hz), results indicated a significant main effect of the lowest cut-off (F(_3,20_)=8.816, p< 0.001) but no effect of stimuli type (F(_1,20_)=0.272, p=0.608) and no significant interaction (F(_3,60_)=2.127, p=0.106).

**Fig. 3.**
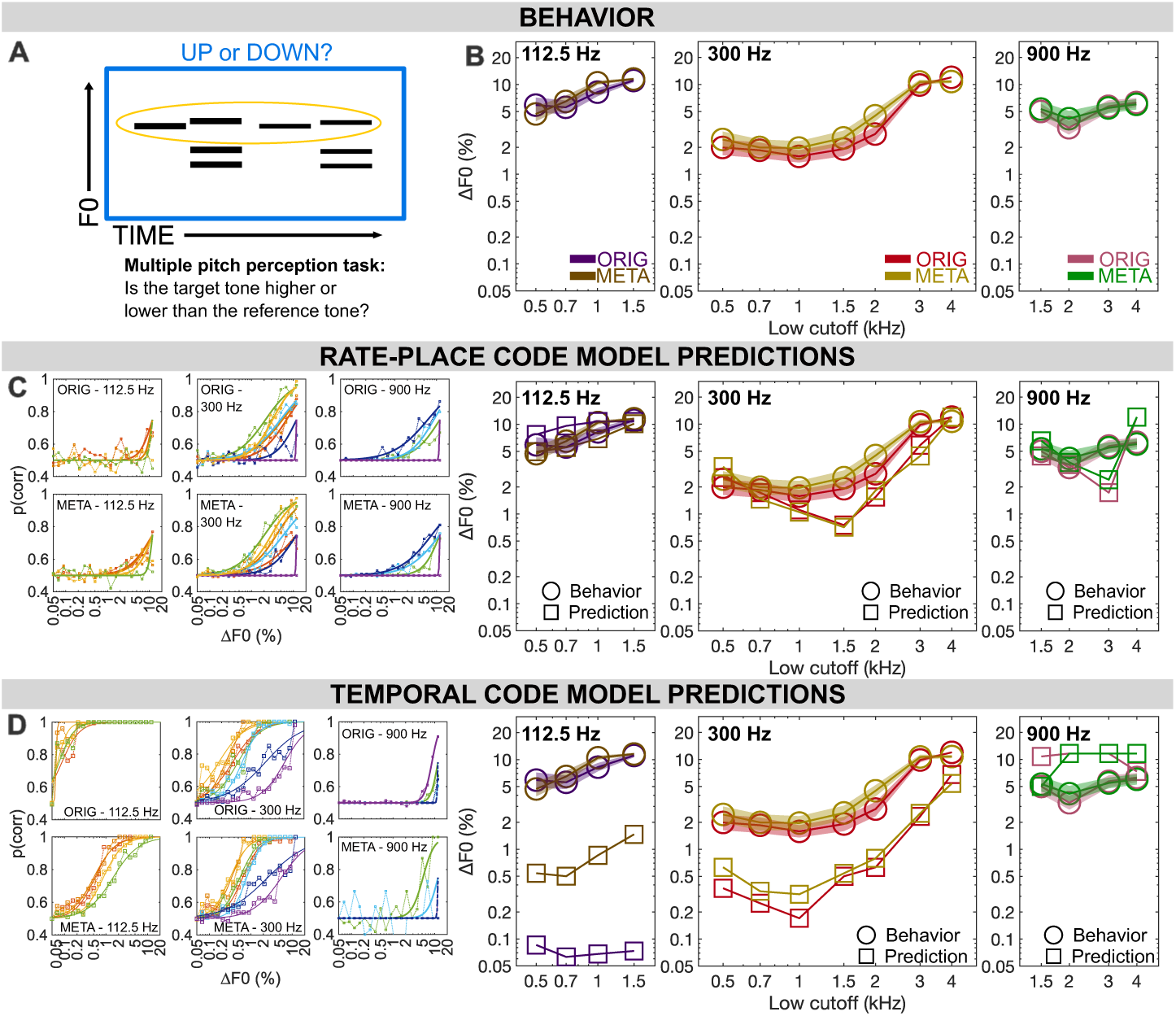
Results from Experiment 2: Multiple pitch discrimination. **(A)** Participants were asked to judge the direction of pitch change between notes in the top voice of the illustrated multi-voice stimulus, ignoring the lower two voices. In one condition (ORIG), each note was a harmonic complex tone in noise. In the other (META), each sound was a rate-place metamer of a single pitch or mixture of pitches. **(B)** F0 difference limens by spectral region, tone type, and F0. **(C)** Predictions from a rate-place model, “with neurometric functions by condition on the left and corresponding threshold predictions on the right, plotted against human behavioral thresholds. **(D)** Predictions from a temporal model, as in (C) but using summary autocorrelation functions rather than rate-place profiles. Throughout: shaded regions ± 1 SEM.

Behavioral results for Experiment 2 overall indicate that pitch discrimination in the context of multiple simultaneous pitches is surprisingly comparable for META and ORIG stimuli. This was true even in frequency regions where listeners only would have access to residual rate-place cues (for e.g. F0=300 Hz, lowest cutoff 1.5 kHz), where there may be identifiable peaks and troughs in the rate-place profile, but without obvious correspondence to individual harmonics of any of the three F0s. Figures 3C and 3D show behavioral thresholds predicted by models of pitch discrimination using rate-place and autocorrelation cues respectively. As in Experiment 1, the autocorrelation model significantly overestimates precision for resolved stimuli, while failing to complete the task at all in fully unresolved stimuli, resulting in a predicting a large gap between ORIG and META. While the rate-place predictions do not provide a perfect match to human behavioral data, they come much closer than the autocorrelation model, as demonstrated again by comparison of RMSe. In experiment 2, the rate-place model had an average RMSe across conditions of 1.266, while the temporal model had an average RMSe across conditions of 2.472 (see also Table 2 in Supplemental Methods).

### Rate-place cues are likely more important for multi-pitch masked pitch discrimination

The rate-place model provided overall a close match to the behavioral results for Experiment 2. The temporal model predicts an enormous gap between ORIG and META for low-F0 unresolved pitch, with overestimation of performance for ORIG stimuli. This is not observed behaviorally, suggesting that human listeners are unlikely to rely on these cues for this task. The temporal model also predicts a steep shift from resolved to unresolved spectral regions for mid-range F0s, which differs from the more gradual shift in behaviorally measured thresholds.

Overall, comparison of modeled rate-place and timing cues for hearing out a single pitch within a mixture of HCTs reveals likely dominance of rate-place cues, albeit with some evidence that the rate-place model alone may not represent the only information integrated in these decisions. Modeled timing cues, unlike rate-place cues, are severely degraded by the rate-place metamerization process. This degradation, however, leads to no corresponding change in human discrimination thresholds. Slight differences in ORIG and META behavioral results still lack a clear explanation, but reliance on timing cues seems an unlikely one.

### Experiment 3: Discriminating pitch mixtures in musically relevant tasks

Experiment 2 asked listeners to discriminate a target pitch in a mixture, an ability which relates to ecologically valid tasks such as hearing out a speaker of interest in a group conversation or a melody of interest against a complex background. However, the goal of music perception is not always to hear each note or instrument separately, but also to perceive how constituent parts relate to a greater whole. In the context of musical pitch, certain pitch combinations create harmonies that can be perceived as a gestalt. In Experiment 3, we asked whether rate-place cues could explain human performance in a task designed to index this higher-order processing of the relationship between simultaneous pitches. Listeners (n = 20) were presented with a chord on every trial and asked to label its quality, either “major” or “minor.” To determine whether a chord is major or minor, participants must perceive all 3 F0s in the mixture to a precision of at least 1 semitone, as a 1-semitone error can change any minor triad to a major triad, or vice-versa.

### Rate-place metamers of pitch mixtures are less discriminable than original mixtures in some conditions

Behavioral results for Experiment 3 are shown in Figure 4B in terms of the sensitivity index d’, a measure of discrimination accuracy. Similar to Experiment 2, we performed a repeated measures ANOVA for each spectral region separately to compare the effect of stimulus type (ORIG vs META) and lowest frequency cut-off for each of the 3 F0 conditions. For the low frequency conditions (F0 = 112.5 Hz), we found a significant main effect of stimulus type (F(_1,19_)=8.641, p=0.008) as well as a significant main effect of lowest harmonic cut off (F(_3,57_)=3.031, p=0.037) but no significant interaction (F(_3,57_)=0.485, p=0.694). For the mid frequency conditions (F0 = 300 Hz), a significant main effect was found for stimulus type (F(_1,19_)=29.857, p<.001) as well as lowest harmonic cut off (F(_6,114_)=25.824, p<.001) but no interaction (F(_6,114_)=1.692, p=0.129). For the high frequency conditions (F0 = 900 Hz), no significant main effect of stimulus type was found (F(_1,19_)=2.404, p=0.138) but results indicate a significant main effect of lowest harmonic cut off (F(_3,57_)=13.008, p<0.001) but no significant interaction (F(_3,57_)=0.898, p=0.448).

**Fig. 4.**
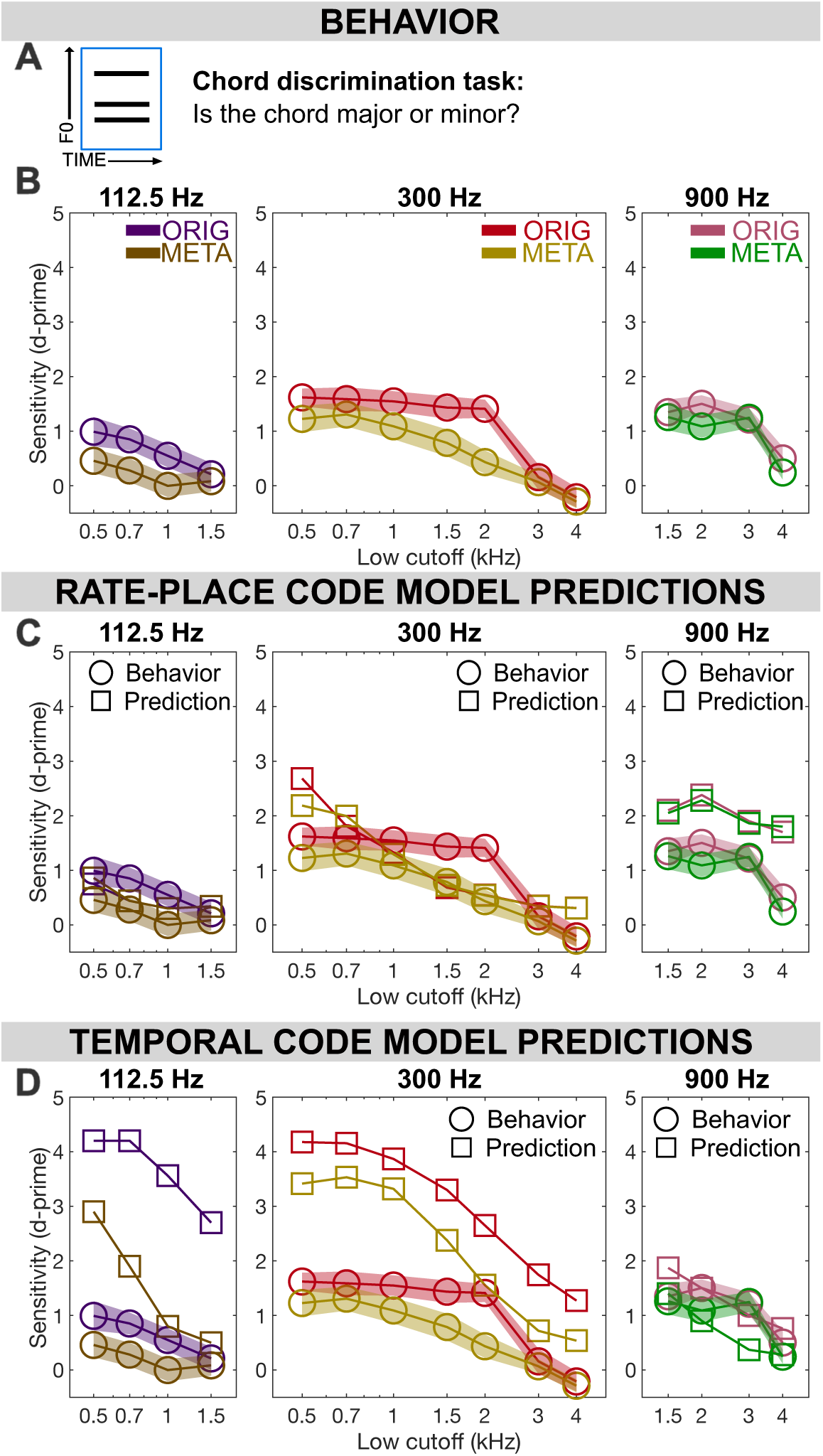
Results from Experiment 3: Chord discrimination. **(A)** Participants were asked to label a three-pitch mixture as either a major or minor triad, with the mixture presented either as harmonic complex tones in noise (ORIG) or a rate-place metamer of the mixture (META). **(B)** Chord discrimination sensitivity (d’) is plotted as a function of spectral region, tone type, and F0. **(C)** Predicted sensitivity from a rate-place model in each condition, plotted against behavioral results from humans. **(D)** Predictions from a temporal model, as in (C) but using summary autocorrelation functions rather than rate-place profiles. Throughout: shaded regions ± 1 SEM.

### Integration of rate-place and timing cues may be important for discriminating pitch mixtures

Human behavioral data (Figure 4B) shows a significant gap in performance between ORIG and META in unresolved conditions, including mid-F0 high-frequency conditions as well as low-F0 conditions in general. This gap is not accounted for by the rate-place model, suggesting integration of timing cues is necessary to achieve human performance for ORIG stimuli. However, the timing model alone also fails to account for many features in the behavioral data, with global overestimation of performance and an exaggerated gap between ORIG and META.

When comparing model predictions against human behavioral data for Experiment 3, the rate-place model again outperformed the autocorrelation model. Specifically, the RMSe of model predictions for the rate-place model across all conditions was 1.054 in d’ units. Meanwhile, the RMSe across conditions for the temporal model was 1.774 in d’ units (see Table 3 in supplemental methods for RMSe details).

Nevertheless, the gap between ORIG and META is not predicted or explained by the rate-place model, suggesting that timing information is somehow integrated in this task. This finding reveals an important potential difference in neural processing strategies for hearing an isolated pitch in a mixture as opposed to characterizing the mixture itself, especially since the mixtures of HCTs used for both Experiments 2 and 3 were identical. While rate-place representations were sufficient to support pitch perception with concurrent maskers in Experiment 2, significant gaps emerged for harmony perception in Experiment 3.

### Experiment 4: buildup of expectation for pitch mixtures in musical context

Experiments 1-3 measured pitch behaviors ranging in complexity, but all the stimuli presented lacked temporal context. Listeners perceive components of music, such as chords, relative to recently established context, such as a chord progression that establishes what chord to expect next. Experiment 4 sought to determine whether rate-place information, beyond mere major vs. minor distinction for chords in isolation, can support harmony identification in a more ecologically valid context where the chord is evaluated relative to its recent preceding context.

In Experiment 4 (n=20), listeners heard sequences of 7 chords and were asked to rate how expected or unexpected the final (target) chord was, given its context. The buildup of expectation in music is the main feature that allows participants to derive pleasure from it, as reward sensations are linked to the formation of expectations and the confirmation or violation of these expectations (31). The pleasure derived from these expectations stem from two distinct sources: sensory and cognitive expectations (32–36). Sensory expectations are generated by shared acoustic features, such as overlapping harmonic spectra, repetitions of notes in a sequence, or similar timbres. In contrast, cognitive expectations arise from the listener’s accumulated knowledge of musical patterns and structures, acquired through long-term exposure to music.

Although previous experiments explored various F0 ranges, we limited Experiment 4 to a middle F0 range (centered on 300 Hz) to focus on the transition from resolved to unresolved for these F0s. As in previous experiments, we tested both ORIG and META conditions. We also varied the amount of sensory priming in the context preceding the target tone where the conditions without pitch priming ensured that the notes in the final target chord were not present in any of the context chords.

#### Pitch mixtures are identifiable in musical context with rate-place metamers

Behavioral ratings shown in Figure 5B show that listeners consistently were able to rate expected and unexpected chords in the sequence with accuracy for both ORIG and META stimuli, even as rate-place cues were degraded by increasing the lower cutoff of the stimulus bandpass filter to eliminate low-order resolved harmonic, supporting previous findings in multiple pitch perception tasks. For the highest frequency range where rate-place cues would be severely degraded, we see that performance is affected for both ORIG and META stimuli.

**Fig. 5.**
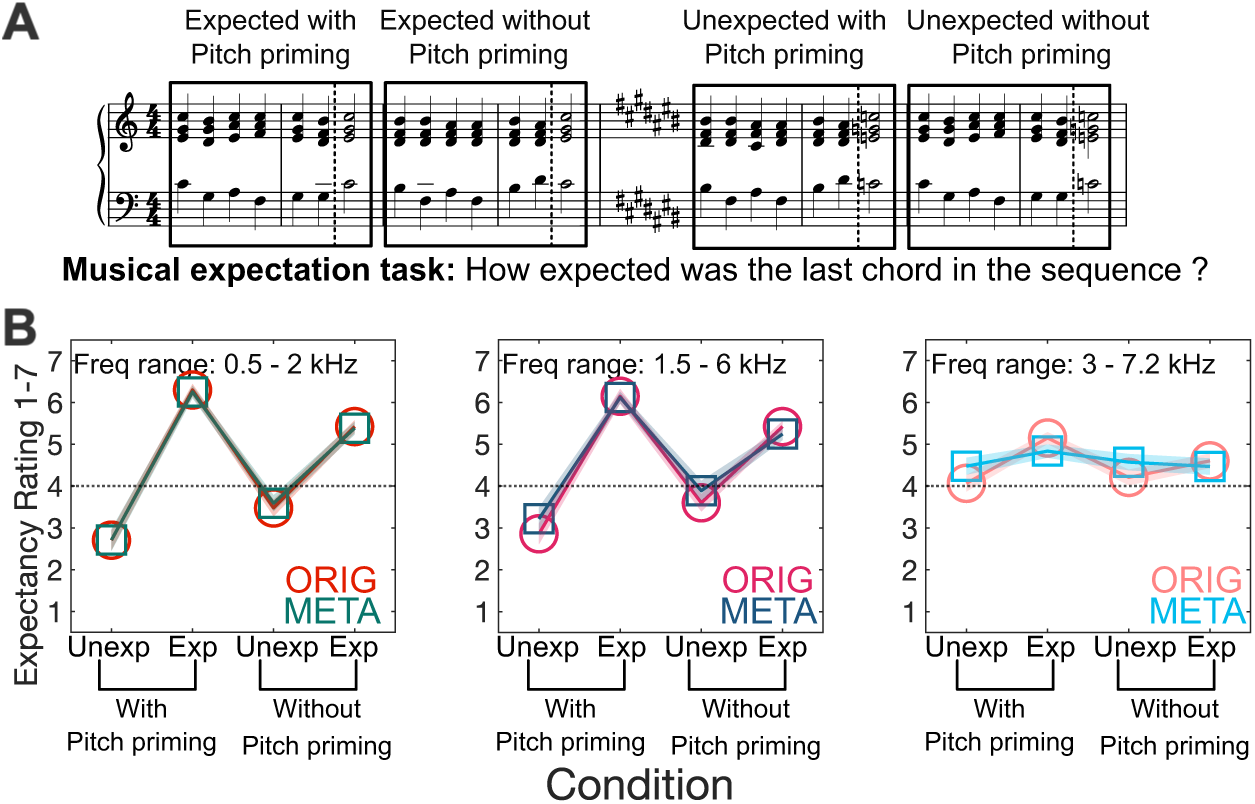
Results from Experiment 4: Musical expectation. **(A)** Participants were asked to rate the degree of expectedness of the final chord in a seven-chord sequence, with sequences designed to elicit strongly divergent expected and unexpected ratings, either including shared pitches between context and target (“with pitch priming”) or not (“without pitch priming”). Each chord was presented either as a mixture of harmonic complex tones in noise (ORIG) or a rate-place metamer of this mixture (META). **(B)** Expectancy ratings by condition for both ORIG and META, with shaded regions ± 1 SEM.

A repeated measures ANOVA with 3 factors was run to compare the effect of stimulus type (ORIG vs META), frequency ranges and sensory priming. A significant main effect was found for stimulus type (F(_1,19_)=49.839, p<0.001), frequency range (F(_2,38_)=65.492, p<0.001) as well as sensory priming (F(_1,19_)=98.066, p<0.001). A significant interaction was found between frequency range and sensory priming (F(_2,38_)=36.317, p<0.001) but no significant interaction of stimulus type and sensory priming (F(_2,38_)=0.040, p=0.844).The three-way interaction between stimulus type, frequency range and sensory priming (F(_2,38_)=0.997, p=0.379) was also not significant.

Overall, rate-place cues supported pitch perception of single complex tones, whether in isolation (Experiment 1) or with concurrent maskers (Experiment 2). However, more complex and ecologically relevant tasks (Experiments 3 and 4) requiring true harmony perception revealed differences between ORIG and META stimuli, suggesting the potential integration of neural timing cues, though the rate-place model still outperformed the timing model for Experiment 3.

## Discussion

We observed that stimuli preserving rate-place representations of pitch also preserve human behavioral responses on pitch tasks in most, but not all, situations. Results for ORIG largely replicate past explorations of the limits of pitch perception (2, 28, 37, 38), validating the comparison with META stimuli from an established baseline. We also expanded the range of F0-related tasks to include not only traditional pitch discrimination threshold measurements, but also more complex musically relevant tasks with multiple pitches. In this way, our results bridge the gap from the limited psychoacoustic literature on multiple pitch perception (20, 21, 39–41) to studies of harmony and harmonic expectation in chord sequences (34–36, 42–44).

Our approach to dissociating potential neural codes for pitch relies on a physiologically grounded model (26) combined with rigorous psychophysics. We address the shortcomings of current models of multiple complex pitch (20, 45, 46) with a closed-loop approach that combines model-based stimulus generation with forward modeling of behavior using the same model of auditory nerve firing.

Overall, we found support for rate-place explanations of pitch perception when listeners focused on isolating a target pitch (Experiments 1-2), in that our rate-place model provided a much closer match to predicting these results than our temporal model. Although the rate-place model continued to perform better overall, we also found some evidence for the importance of timing cues when evaluating chords in harmonic context (Experiments 3-4). This points to potential differences in neural coding at different levels of musical complexity. Our approach constrains the information available at the peripheral stage of processing, not only in modeling but in the stimulus itself. Harmonic expectation (Experiment 4) likely involves higher-level processes that fall beyond the scope of the modeling but are still affected by these peripheral limitations in the stimulus. Therefore, we speculate that temporal representations may be incorporated at higher levels of musically complex pitch processing.

One potential role for temporal representations of multiple-pitch stimuli may come from the perception of harmony as timbre. A quality of sound usually thought of as distinct from pitch but with some overlapping neural representation (47–53), timbre is highly sensitive to timing (54, 55). Indeed, harmony, though it is built from combinations of pitches, has sometimes been studied as an aspect of timbre, given its lack of a clear linear dimension like (single) pitch (56, 57). It seems plausible that the process of integrating multiple pitches into a single timbre-like quality is more likely to incorporate neural timing codes in perceptual decisions, even when these codes are generally not necessary for single pitch perception.

Our modeling approach is not without its pitfalls and assumptions, especially because key model parameters cannot be directly measured in humans. This means our results are dependent on assumptions about tuning, phase locking limits, other aspects of early auditory processing that could be different in other models. Our choice to model low spontaneous-rate fibers at relatively low sound levels was intended to provide the best chance of preserving rate-place cues, as demonstrated in Figure S2. The focus on low spontaneous-rate fibers strengthens the conclusion that rate-place cues were insufficient to convey all information in Experiments 3-4 but may weaken the conclusion that rate-place cues were largely sufficient in Experiments 1-2. Future research could evaluate the effects of varying sound level and include different fiber types in modeling considerations. Also, although rate-place metamers preserve long-term average rate-place profiles, they do not perfectly preserve rate-place cues for every individual stimulus at a trial-by-trial level, which introduces an undesirable difference between META model predictions and META behavioral results. Moreover, it does not completely remove temporal cues in every condition, as at some level place and timing cues are fundamentally intertwined. This admittedly imperfect dissociation between rate-place and timing cues is nevertheless a meaningful advance over previous attempts to resolve the question. This new approach nevertheless significantly advances our ability to dissociate neural cues in intermediate gray zone with “partially resolved harmonics”, where classical approaches are ill-suited to characterize cue fidelity.

Forward models of specific neural codes, such as the approach in the present study, have certain advantages and disadvantages compared to machine-learning strategies relying on deep neural networks (24, 25, 58, 59). DNNs can generally be trained to achieve human-like performance given training inputs that match the natural sound environment of humans. However, the inner mechanisms of neural computation that underlie this performance can remain mysterious beyond a broad understanding of input-output relationships and are unlikely to mimic human cognition on an algorithmic level. Our approach, by contrast, often shows stark differences between model predictions and real human behavior, but these differences can be instructive when produced by models that are explicit, transparent, and describable in physiologically valid terms. This class of model can be modified and interrogated at various levels of processing. Perhaps future research may seek to integrate the two approaches and extract the benefit of each, testing explicit assumptions about neural codes where possible while bridging gaps with machine-learning strategies where necessary. With careful development, such an approach may gradually approximate a human-like auditory system on both algorithmic and performance levels.

## Materials and Methods

### Listeners

In total, 30 listeners (18 female, 10 male, 2 nonbinary) participated in at least one of Experiments 1-3. Participants ranged in age from 19 to 28 years old (mean = 25.5) and had formal music training ranging from 0 to 18 years (mean = 2.3). These 30 participants were trained on pure-tone versions of Experiments 1-3 (see “Procedure” for these Experiments below) to determine eligibility to participate in any of these experiments. From this group of 30 participants, 10 failed to pass screening for at least one of Experiments 1-2, disqualifying them from both experiments, and thus only completed Experiment 3. Another 10 failed to pass screening for Experiment 3, and thus only completed Experiments 1-2. The remaining 10 completed all 3 experiments.

Participants in Experiments 1-3 were screened for hearing thresholds at 14 kHz using a 2-alternative forced-choice adaptive tracking procedure for detection of a tone in noise. The cutoff for inclusion was a threshold under 40 dB SPL for tones presented with threshold-equalizing noise (TEN) at 35 dB SPL in the equivalent rectangular bandwidth (ERB) centered at 1 kHz. Of the 29 participants included, 12 passed the 14 kHz screening task. Participants who failed the high-frequency screening were excluded from the high-frequency conditions of Experiment 1 but were included in analyses for the remainder of the study, which did not include task-relevant sounds above 8 kHz.

A new group of 20 normal-hearing listeners (11 female, 6 male, 3 nonbinary) was recruited to participate in Experiment 4. These participants ranged in age from 18 to 30 years old (mean = 23.4 years) and had formal music training ranging from 0 to 17 years (mean = 4.7). As in Experiments 1-3, these participants were also tested for audiometric thresholds at or below 20 dB HL at octave frequencies from 0.25 – 8 kHz. However, for Experiment 4, no other performance criterion was applied, and all 20 participants who passed audiometric testing went on to complete Experiment 4.

All participants in all experiments provided informed consent before participation and were compensated at a rate of $15 per hour. Study procedures were approved by the University of Michigan Medical School Institutional Review Board (IRBMed HUM00218552).

### Stimuli

Stimuli in all 4 experiments included original pitch-evoking stimuli (ORIG) and rate-place metamers of those stimuli (META). All ORIG stimuli in Experiments 1-4 were bandpass-filtered harmonic complex tones (HCTs) with harmonic components at 45 dB SPL within the passband, masked by broadband TEN (60) at a level of 35 dB SPL within the equivalent rectangular bandwidth (ERB) at 1 kHz (61). Additional TEN at 40 dB SPL per ERB was also included at low frequencies, with a high cutoff 1 kHz below the stimulus passband, to ensure the F0 region was well masked. The bandpass filters applied to HCTs in ORIG stimuli were constructed slightly differently in Experiment 1 than in Experiments 2-4. In Experiment 1, bandpass filters were applied as in (28), with a low cutoff equal to the 4th, 5th, 7th, 8th, 10th, 11th, 13th, or 14th multiple of the F0, and a high cutoff 9 ranks higher (e.g., at the 13th component when the low cutoff was the 4th). Components outside the passband decreased in level by 6 dB per rank. In Experiments 2-4, as in (45), the passband did not depend on F0 but remained constant across a given condition. From low to high, the seven spectral regions tested in Experiments 2-4 were: 0.5-2, 0.7-3, 1-4, 1.5-6, 2-6.6, 3-7.2, and 4-8 kHz. Components outside the passband decreased in level by 0.03 dB per Hz (equivalent to 6 dB every 200 Hz).

For every ORIG stimulus in the main experiment, we created its META equivalent by iteratively filtering samples of Gaussian noise (see Figure 1A for a schematic illustration of this process), loosely based on model-metamer strategies of previous studies with visual and auditory metamers (58, 62). We first filtered a sample of Gaussian noise using the same filter parameters as the passband of the ORIG stimulus, with a flat spectrum in the passband and slopes of 6 dB/rank of the corresponding F0 (Experiment 1) or 0.03 dB/Hz (Experiments 2-4). We added TEN as in the ORIG condition, creating a noise stimulus with the same broad spectral shape as the ORIG stimulus. We then simulated the rate-place representation of the ORIG stimulus and the noise sample using an established model simulating firing at the level of the auditory nerve (26). The model responses were generated using low-spontaneous-rate fibers in the humanized variant of the model. Fractional Gaussian noise parameters were set to variable, and the model was set to “spike” mode. For both the noise stimulus and the target stimulus, we simulated an auditory nerve average rate (ANAR) response by averaging the number of spikes over time for 10 samples of 1000 ms each, discarding the first and last 100 ms of each response. The ANAR was simulated for a range of 200 characteristic frequencies (CFs), logarithmically spaced, from 300 Hz below the low-frequency cutoff to 300 Hz above the high-frequency cutoff for each stimulus. By subtracting the ANAR of the target ORIG stimulus from the ANAR of the noise stimulus, we obtained a spectral difference function, which was subsequently used to filter the noise in the spectral domain in order to reduce the difference between the ORIG ANAR and noise sample ANAR.

On each iteration of the metamerization process, we adjusted the magnitude spectrum of the noise stimulus to bring its ANAR closer to the target ANAR, using an incremental learning rate (0.1 dB SPL per spike/s), akin to the metamerization method of (62). The iterative adjustment process is necessary due to nonlinear properties of the model response. Fifty iterations were sufficient to match the ANAR profiles within 1 spike/s across cochlear channels. Figure 1B illustrates the strong correspondence in ANARs and clear spectral differences between ORIG and META stimuli, showing that, while their acoustic spectra differ, their rate-place representations are matched.

### Procedure

In brief, the study was divided into 4 experiments, each of which measured different aspects of pitch perception or multiple pitch perception. After audiometric screening, participants in each experiment were given verbal instructions and practice with a training version (5 training blocks of 20 trials each) of that experiment, using pure tones with no masking noise. A participant was determined to have passed the training procedure if they achieved a sensitivity index (d’) of 1.5 or higher on any of the training blocks for that experiment.

Experiment 1 measured F0DLs for single HCTs. The participants were asked to report whether the second tone was higher or lower in pitch than the first. ΔF0 between the two tones was adaptively varied using a 1-up 2-down rule until the sixth reversal at final step size, with each threshold the average of 2 runs per condition in a randomized order. The maximum ΔF0 allowed in this procedure was 2 semitones.

Experiment 2 measured F0DLs for a single pitch presented within a mixture of three simultaneous HCTs with three different F0s. On each trial, participants heard four sounds. Sounds 1 and 3 were single HCTs while sounds 2 and 4 were triads (mixtures of 3 HCTs) in which the tone with the highest pitch was close in F0 to the F0 of sounds 1 and 3. Participants were asked to compare the reference pitch in sounds 1 and 3 to the target highest pitch in sounds 2 and 4, and say whether the pattern of pitch changes was upwards or downwards. Thresholds were averaged across 3 runs per condition in a randomized order. As in experiment 1, the maximum ΔF0 allowed in this procedure was 2 semitones.

Experiment 3 was adapted from (45) and measured participants’ accuracy at discriminating different types of triads, requiring participants to identify all three F0s in the mixture to within 1 semitone precision. On each trial, participants heard a single triad and were asked to assign it as either Major or Minor.

Experiment 4 asked participants to give subjective ratings of expectedness for the final chord in a sequence of 7 chords. Participants provided this rating on a scale from 1-7, where 1 was “least expected” and 7 was “most expected.”

### Modeling

Auditory nerve model responses were computed in response to all stimuli used experiments 1-3, and “F0 salience functions” were computed using two different strategies. One model extracts only rate-place cues, ignoring neural response timing. The other model is based on extracting temporal cues, ignoring average firing rates. Discrimination performance was then estimated from differences in F0 salience functions in each of these two models. For Experiment 1, neurometric functions were computed to estimate the sensitivity index d’ at a range of values for ΔF0, with the predicted threshold corresponding to the point where d’ crosses 1. For Experiment 2, neurometric functions were computed for percent correct (pcorr) at a range of ΔF0, with predicted threshold where pcorr crosses 70%. For Experiment 3, the sensitivity index d’ was directly estimated by the model and compared against performance in humans.

### Experiment 1

#### Stimulus preparation

Simulated auditory nerve responses were computed in response to a total of 38400 stimuli, divided into 19200 ORIG and 19200 equivalent META. Each category contained 8 examples each for 20 simulated model “trials”, at 20 different ΔF0 values, logarithmically spaced between 0.05% and 12% ΔF0, for a total of 3200 stimuli in each of the 6 frequency conditions. The 8 examples were divided along three different factors: high or low F0 (separated by a distance of ΔF0), lowest harmonic rank (depending on condition, e.g. 4^th^ vs. 5^th^ or 7^th^ vs. 8^th^), and first or last half of the 1000-ms stimuli. By computing separate responses for the first and last half of the 1000 ms pre-generated metamer stimuli, we created 500-ms stimuli equivalent to those heard by humans during the experiment.

##### Auditory nerve model response

Each run of the auditory nerve model simulated 200 different characteristic frequencies (CFs), logarithmically spaced between 150 Hz and 9600 Hz. These simulations used low spontaneous-rate fibers, tuning parameters optimized for normal-hearing humans, and variable fractional Gaussian noise associated with spontaneous activity of AN fibers. The stimulus was resampled at 100 kHz for modeling purposes and scaled to the appropriate level relative to the reference pressure level of 20 x 10^-6^ pascals.

##### Rate-place feature extraction

This model extracts only rate-place cues, ignoring neural response timing. The auditory nerve average rate (ANAR) was extracted from each model response by averaging the responses across time in each channel over the period from 100ms to 500 ms post stimulus onset. In order to extract a feature directly linked to F0 from the ANARs, we computed a fast Fourier transform (FFT) of ANARs along the frequency axis, creating a response we called FFT-ANAR. Rates for channels outside the passband were set to 0 before computing the FFT-ANAR, to avoid including artifacts related to edge frequencies. The FFT-ANAR contains peaks corresponding to F0s, without incorporating any information about neural response timing.

##### Autocorrelation feature extraction

This model extracts only neural response timing cues, ignoring rate-place cues. The autocorrelation was computed in each channel at a range of 200 lags from 0.3 to 19.2 ms, corresponding to F0s from 52 Hz to 3333 Hz. These autocorrelation functions were then summed across channels, creating summary autocorrelation functions (SACFs). The resulting SACFs contain peaks corresponding to F0s in the original stimuli, and can be used to model pitch task behavior in an equivalent manner to the FFT-ANAR signals in the rate-place model.

##### Estimation of sensitivity d’

From this point forward in the modeling pipeline, FFT-ANARs and SACFs are handled in equivalent ways, both referred to now as F0SF (F0 strength functions). The modeling procedure resulted in 8 F0SFs for each modeling “trial.” These F0SFs were then compared to each other on a pairwise basis, with each of the 4 high-F0 stimuli crossed with each of the 4 low-F0 stimuli to evaluate the “target distribution” of F0SF distances. The distances were computed as Euclidean distances between F0SFs. For comparison, the “reference distribution” of F0SF distances contained all the pairwise distances between F0SFs in response to stimuli with the same F0. The estimated sensitivity d’ was then taken as the difference between target and reference distributions, normalized by their pooled standard deviation.

##### Neurometric functions and threshold estimates

In every condition, as ΔF0 increases, the resulting estimated sensitivity d’ also increases, starting from the chance level of 0 and gradually increasing to a ceiling value. This pattern of gradually increasing sensitivity is the neurometric function. See left panels of Figure 2C and 2D for modeled neurometric functions in each condition of Experiment 1. We artificially limited d’ to 2, since we sought to estimate thresholds where d’ = 1. We estimated thresholds by fitting a sigmoid function to the neurometric function in each condition, with only 2 parameters varying during the sigmoid fitting process: steepness k and threshold x0. The floor of the sigmoid was fixed at 0 and the ceiling was fixed at 2. The point where this fitted sigmoid crosses 1 was then taken as the estimate of the threshold for that condition. At the end of the process, the thresholds can be reasonably compared against human behavioral thresholds for estimates of possible performance using each version of the model

#### Experiment 2

In order to establish model predictions for experiment 2, as in experiment 1, we presented stimuli to the auditory nerve model equivalent to those presented in the human behavioral experiment. For the ORIG condition, stimuli were harmonic complex tones as well as triadic mixtures of three simultaneous HCTs. Each HCT or triad was filtered into a different frequency region (replicating each of the 15 different conditions used in the behavioral experiment, each covering 1-2 octaves between 500 Hz and 8000 Hz) and with a different range of F0s (centered around either 112.5, 300, or 900 Hz). The level per component was set to 45 dB SPL. ORIG stimuli also contained background TEN set to 35 dB SPL per ERB. An additional masking noise at 30 dB/ERB was added, lowpass filter to 1.5 components below the lowest component in the included mixture. As in Experiment 1, META stimuli were metamers of ORIG stimuli, drawn from pre-computed metamers of a range of HCTs and triads in the same F0 and frequency conditions.

##### Stimulus preparation

Model responses for experiment 2 were calculated for a total of 96000 stimuli, divided into 48000 ORIG and 48000 equivalent META. In each condition, we ran 800 triads (crossing 2 exemplars, 20 ΔF0 values, and 20 trials) through the model, as well as 2400 single HCTs (crossing 3 F0 values, 2 exemplars, 20 ΔF0 values, and 20 trials), for a total of 3200 stimuli per condition for each of ORIG and META. The ΔF0 values as in experiment 1 were logarithmically spaced between 0.05% and 12% ΔF0. As in experiment 1, the 2 exemplars were either the first and last half of the pre-generated 1000-ms META stimulus, or a newly generated 500-ms ORIG stimulus. The 3 F0 values for the single tones were either identical to the target (highest) F0 in the mixture, or up or down by ΔF0.

##### Auditory nerve model responses and F0 feature extraction

These were collected for each stimulus in the same way as in Experiment 1 and passed through rate-place and autocorrelation-based feature extraction in the same way.

##### Estimation of percent correct, neurometric functions and threshold estimates

Each modeling trial consisted of 6 F0SFs for single tones and 2 F0SFs for triads. These F0SFs were then compared on a pairwise basis, simulating the behavioral paradigm comparing reference tones to target notes in the triad. The model’s choice was evaluated as correct or incorrect on each trial, producing neurometric functions which were then used as in Experiment 1 to produce estimates of discrimination thresholds.

#### Experiment 3

In the modeling strategy for Experiment 3, we followed the same procedures for stimulus preparation as for the triads in Experiment 2, excluding the single HCT stimuli as they were not relevant to Experiment 3. Instead of estimating F0s within triads, however, we now directly compared F0SFs for major triads to minor triads, estimating d’ for the major-minor distinction as the equivalent of a single point on the neurometric function (similar to the constant-stimulus paradigm employed for humans on this task). Resulting d’ was compared to d’ measured directly in humans, without estimating thresholds.

#### Experiment 4

Due to the complex nature of the behavioral task completed by listeners in Experiment 4, direct behavioral predictions were beyond the scope of this paper. However, we note a limiting factor in performance on this task: differentiating “expected” from “unexpected” stimuli in ratings requires listeners to discriminate between chords separated by one semitone in all three F0s. Modeled performance on Experiment 3, which requires discrimination between chords that differ by one semitone in only one F0, can thus be taken as a ceiling for any model of Experiment 4. Behavioral results for Experiment 4 were evaluated with this in mind.

## Acknowledgments

This study was supported by NIH grant R00 DC017472 (AHM). We thank Dr. Laurel Carney for advice on the computational modelling.

## Data Availability

Behavioral data and code from all experiments in this article have been deposited in Open Science Framework: https://osf.io/hkyqm/overview?view_only=825d66afe879462c93287b8d82f1a3b7

## Notes

https://osf.io/hkyqm/overview?view_only=825d66afe879462c93287b8d82f1a3b7

### Competing Interest Statement

The authors have declared no competing interest.

## References

1. H. von Helmholtz, A. J. Ellis, On the sensations of tone as a physiological basis for the theory of music (London, New York : Longmans, Green, and Co., 1895).

2. A. Houtsma, J. Smurzynski, Pitch identification and discrimination for complex tones with many harmonics. The Journal of the Acoustical Society of America 87, 304–310 (1990).

3. J. C. R. Licklider, A duplex theory of pitch perception. Experientia 7, 128–134 (1951).

4. P. A. Cariani, B. Delgutte, Neural correlates of the pitch of complex tones. I. Pitch and pitch salience. Journal of Neurophysiology 76, 1698–1716 (1996).

5. R. Plomp, Pitch of Complex Tones. The Journal of the Acoustical Society of America 41, 1526 (1967).

6. A. W. Bronkhorst, The cocktail-party problem revisited: early processing and selection of multi-talker speech. Atten Percept Psychophys 77, 1465–1487 (2015).

7. M. Chatterjee, et al., Voice emotion recognition by cochlear-implanted children and their normally-hearing peers. Hearing Research 322, 151–162 (2015).

8. W. A. Yost, Pitch perception. Attention, Perception, & Psychophysics 71, 1701–1715 (2009).

9. A. J. Oxenham, How we hear: the perception and neural coding of sound. Annual Review of Psychology 69, 27–50 (2018).

10. X. Wang, Cortical coding of auditory features. Annual Review of Neuroscience 41, 527–552 (2018).

11. K. M. M. Walker, J. K. Bizley, A. J. King, J. W. H. Schnupp, Cortical encoding of pitch: Recent results and open questions. Hear Res 271, 74–87 (2011).

12. L. Cedolin, B. Delgutte, Pitch of complex tones: Rate-place and interspike interval representations in the auditory nerve. Journal of Neurophysiology 94, 347–362 (2005).

13. Y. Su, B. Delgutte, Robust rate-place coding of resolved components in harmonic and inharmonic complex tones in auditory midbrain. J. Neurosci. 40, 2080–2093 (2020).

14. F. L. Wightman, The pattern-transformation model of pitch. Journal of the Acoustical Society of America 54, 407–416 (1973).

15. M. A. Cohen, S. Grossberg, L. L. Wyse, A spectral network model of pitch perception. The Journal of the Acoustical Society of America 98, 862–879 (1995).

16. R. Meddis, L. O’Mard, A unitary model of pitch perception. J. Acoust. Soc. Am. 102, 1811–1820 (1997).

17. L. Cedolin, B. Delgutte, Spatiotemporal representation of the pitch of harmonic complex tones in the auditory nerve. J. Neurosci. 30, 12712–12724 (2010).

18. S. Shamma, D. Klein, The case of the missing pitch templates: how harmonic templates emerge in the early auditory system. The Journal of the Acoustical Society of America 107, 2631–2644 (2000).

19. J. G. W. Bernstein, A. J. Oxenham, An autocorrelation model with place dependence to account for the effect of harmonic number on fundamental frequency discrimination. The Journal of the Acoustical Society of America 117, 3816–3831 (2005).

20. D. R. Guest, A. J. Oxenham, Human discrimination and modeling of high-frequency complex tones shed light on the neural codes for pitch. PLOS Computational Biology 18, e1009889 (2022).

21. J. E. Graves, A. J. Oxenham, Pitch discrimination with mixtures of three concurrent harmonic complexes. The Journal of the Acoustical Society of America 145, 2072–2083 (2019).

22. J. L. Goldstein, An optimum processor theory for the central formation of the pitch of complex tones. J. Acoust. Soc. Am. 54, 1496–1516 (1973).

23. M. G. Heinz, H. S. Colburn, L. H. Carney, Evaluating auditory performance limits: I. one-parameter discrimination using a computational model for the auditory nerve. Neural Computation 13, 2273–2316 (2001).

24. M. R. Saddler, R. Gonzalez, J. H. McDermott, Deep neural network models reveal interplay of peripheral coding and stimulus statistics in pitch perception. Nat Commun 12, 7278 (2021).

25. M. R. Saddler, J. H. McDermott, Models optimized for real-world tasks reveal the task-dependent necessity of precise temporal coding in hearing. Nat Commun 15, 10590 (2024).

26. M. Zilany, I. Bruce, L. Carney, Updated parameters and expanded simulation options for a model of the auditory periphery. The Journal of the Acoustical … 135, 283–6 (2014).

27. C. Micheyl, M. V. Keebler, A. J. Oxenham, Pitch perception for mixtures of spectrally overlapping harmonic complex tones. The Journal of the Acoustical Society of America 128, 257–69 (2010).

28. A. H. Mehta, A. J. Oxenham, Effect of lowest harmonic rank on fundamental-frequency difference limens varies with fundamental frequency. The Journal of the Acoustical Society of America 147, 2314–2322 (2020).

29. D. R. Guest, A. J. Oxenham, Human discrimination and modeling of highfrequency complex tones shed light on the neural codes for pitch. PLoS Computational Biology 18 (2022).

30. K. L. Whiteford, A. J. Oxenham, Using individual differences to test the role of temporal and place cues in coding frequency modulation. The Journal of the Acoustical Society of America 138, 3093–3104 (2015).

31. D. Huron, Musical aesthetics: Uncertainty and surprise enhance our enjoyment of music. Current Biology 29, R1238–R1240 (2019).

32. V. K. M. Cheung, et al., Cognitive and sensory expectations independently shape musical expectancy and pleasure. Philos Trans R Soc Lond B Biol Sci 379, 20220420 (2024).

33. T. Collins, B. Tillmann, F. S. Barrett, C. Delbé, P. Janata, A combined model of sensory and cognitive representations underlying tonal expectations in music: from audio signals to behavior. Psychol Rev 121, 33–65 (2014).

34. D. R. Sears, M. T. Pearce, J. Spitzer, W. E. Caplin, S. McAdams, Expectations for tonal cadences: Sensory and cognitive priming effects. Quarterly Journal of Experimental Psychology 72, 1422–1438 (2019).

35. B. Tillmann, P. Janata, J. Birk, J. J. Bharucha, Tonal centers and expectancy: Facilitation or inhibition of chords at the top of the harmonic hierarchy? Journal of Experimental Psychology: Human Perception and Performance 34, 1031–1043 (2008).

36. E. Bigand, B. Poulin, B. Tillmann, F. Madurell, D. A. D’Adamo, Sensory versus cognitive components in harmonic priming. J Exp Psychol Hum Percept Perform 29, 159–171 (2003).

37. T. M. Shackleton, R. P. Carlyon, The role of resolved and unresolved harmonics in pitch perception and frequency modulation discrimination. The Journal of the Acoustical Society of America 95, 3529–3540 (1994).

38. E. J. Allen, J. Mesik, K. N. Kay, A. J. Oxenham, Distinct Representations of Tonotopy and Pitch in Human Auditory Cortex. Journal of Neuroscience 42, 416–434 (2022).

39. J. G. W. Bernstein, A. J. Oxenham, Pitch discrimination of diotic and dichotic tone complexes: Harmonic resolvability or harmonic number? The Journal of the Acoustical Society of America 113, 3323–3334 (2003).

40. C. Micheyl, J. G. W. Bernstein, A. J. Oxenham, Detection and F0 discrimination of harmonic complex tones in the presence of competing tones or noise. The Journal of the Acoustical Society of America 120, 1493–1505 (2006).

41. R. T. Penninger, et al., Perception of polyphony with cochlear implants for 2 and 3 simultaneous pitches. Otology & Neurotology 35, 431–6 (2014).

42. S. Leino, E. Brattico, M. Tervaniemi, P. Vuust, Representation of harmony rules in the human brain: Further evidence from event-related potentials. Brain Research 1142, 169–177 (2007).

43. F. Marmel, B. Tillmann, W. Dowling, Tonal expectations influence pitch perception. Perception & Psychophysics 70, 841–852 (2008).

44. S. Koelsch, S. Jentschke, D. Sammler, D. Mietchen, Untangling syntactic and sensory processing: An ERP study of music perception. Psychophysiology 44, 476–490 (2007).

45. J. E. Graves, A. J. Oxenham, Pitch discrimination with mixtures of three concurrent harmonic complexes. The Journal of the Acoustical Society of America 145, 2072–2083 (2019).

46. A. de Cheveigné, H. Kawahara, Multiple period estimation and pitch perception model. Speech Communication 27, 175–185 (1999).

47. E. J. Allen, A. J. Oxenham, Symmetric interactions and interference between pitch and timbre. The Journal of the Acoustical Society of America 135, 1371–9 (2014).

48. E. J. Allen, P. C. Burton, C. A. Olman, A. J. Oxenham, Representations of pitch and timbre variation in human auditory cortex. Journal of Neuroscience 37, 1284–1293 (2017).

49. E. J. Allen, et al., Encoding of natural timbre dimensions in human auditory cortex. Neuroimage 60–70 (2018). 10.1016/j.neuroimage.2017.10.050.Encoding.

50. J. K. Bizley, K. M. M. Walker, B. W. Silverman, A. J. King, J. W. H. Schnupp, Interdependent Encoding of Pitch, Timbre, and Spatial Location in Auditory Cortex. The Journal of Neuroscience 29, 2064–2075 (2009).

51. B. K. Lau, A. J. Oxenham, L. A. Werner, Infant Pitch and Timbre Discrimination in the Presence of Variation in the Other Dimension. JARO - Journal of the Association for Research in Otolaryngology 22, 693–702 (2021).

52. M. J. McPherson, J. H. McDermott, Relative pitch representations and invariance to timbre. Cognition 232 (2023).

53. R. D. Melara, L. E. Marks, Interaction among auditory dimensions: Timbre, pitch, and loudness. Perception & Psychophysics 48, 169–178 (1990).

54. M. D. Hall, J. W. Beauchamp, Clarifying spectral and temporal dimensions of musical instrument timbre. Canadian Acoustics 37, 3–22 (2009).

55. T. R. Agus, C. Suied, S. J. Thorpe, D. Pressnitzer, Fast recognition of musical sounds based on timbre. The Journal of the Acoustical Society of America 131, 4124–4133 (2012).

56. S. McAdams, S. Winsberg, S. Donnadieu, G. De Soete, J. Krimphoff, Perceptual scaling of synthesized musical timbres: Common dimensions, specificities, and latent subject classes. Psychological research 58, 177–192 (1995).

57. B. Tillmann, E. Bigand, N. Escoffier, P. Lalitte, The influence of musical relatedness on timbre discrimination. European Journal of Cognitive Psychology 18, 343–358 (2006).

58. J. Feather, G. Leclerc, A. Mądry, J. H. McDermott, Model metamers reveal divergent invariances between biological and artificial neural networks. Nat Neurosci 26, 2017–2034 (2023).

59. N. Ahmad, I. Higgins, K. M. M. Walker, S. M. Stringer, Harmonic Training and the Formation of Pitch Representation in a Neural Network Model of the Auditory Brain. Frontiers in computational neuroscience 10, 24 (2016).

60. B. C. J. Moore, M. Huss, D. A. Vickers, B. R. Glasberg, J. I. Alcántara, A test for the diagnosis of dead regions in the cochlea. British Journal of Audiology 34, 205–224 (2000).

61. B. R. Glasberg, B. C. J. Moore, Derivation of auditory filter shapes from notched-noise data. Hearing Research 47, 103–138 (1990).

62. J. Freeman, E. P. Simoncelli, Metamers of the ventral stream. Nature neuroscience 14, 1195–201 (2011).

