## Supplemental Figures and Tables for "Dissociating the neural codes for multiple pitch perception in humans"

**Supplementary Information (Figures S1-S4 and Tables 1-3)**

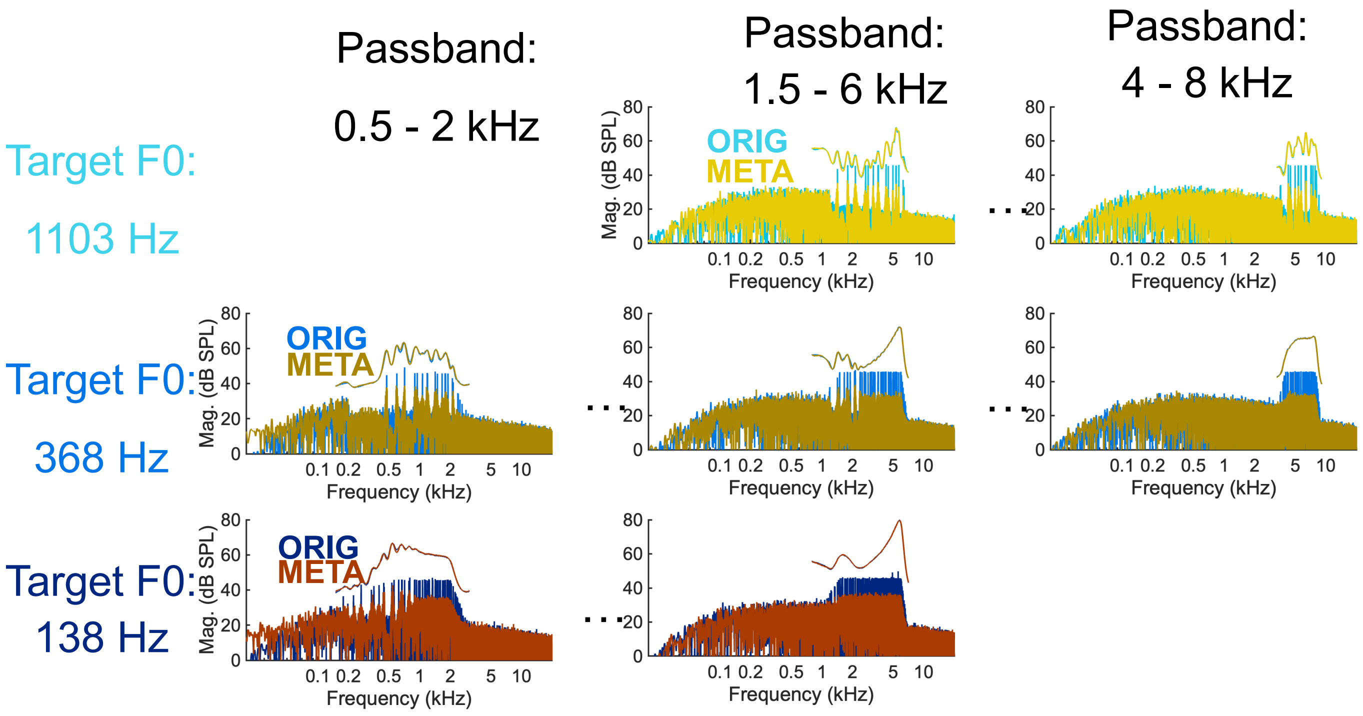


Supplementary Figure 1: Example frequency spectra of original and metamer stimuli at three different F0s and three different spectral regions. Curves above spectra show average firing rate in spikes/s, showing the equivalence in rate-place profile between ORIG and META, despite their acoustically different frequency spectra.

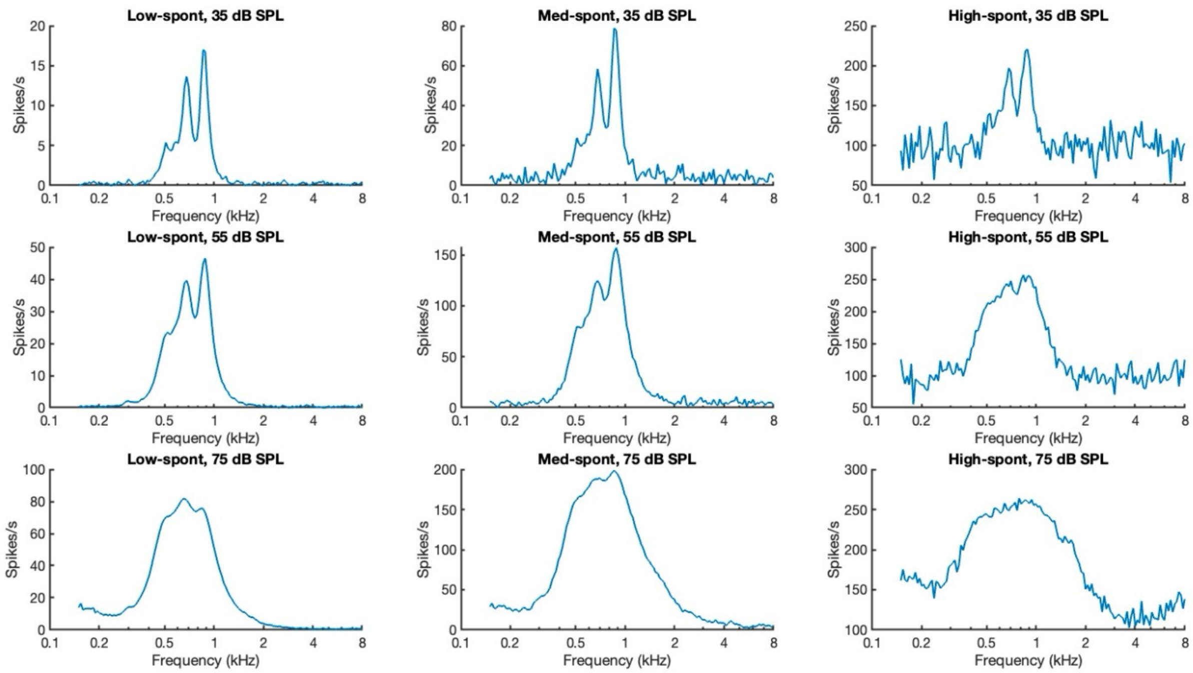


Supplementary Figure 2: Examples of auditory nerve average firing rate responses (ANAR) to a three-F0 mixture filtered between 500 and 1000 Hz, for low, medium and high spontaneous rate fibers at three different sound levels. Because the ANAR shows the clearest peaks with low sound level and low spontaneous-rate fibers, we selected these model and stimulus parameters for all experiments.

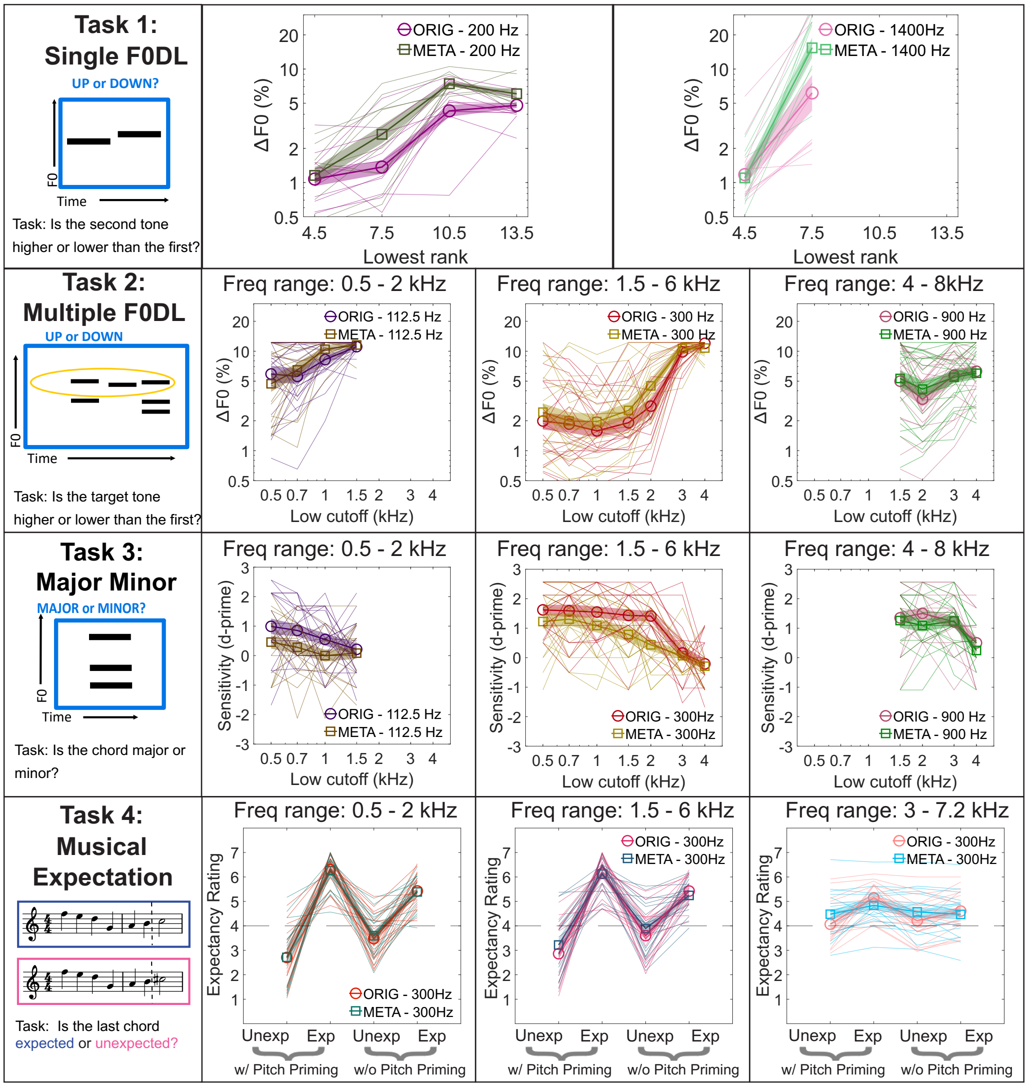


Supplementary Figure 3: Behavioral data plots for experiments 1-4, with individual lines showing each participant's data individually, along with standard error of the mean plotted as shaded regions.

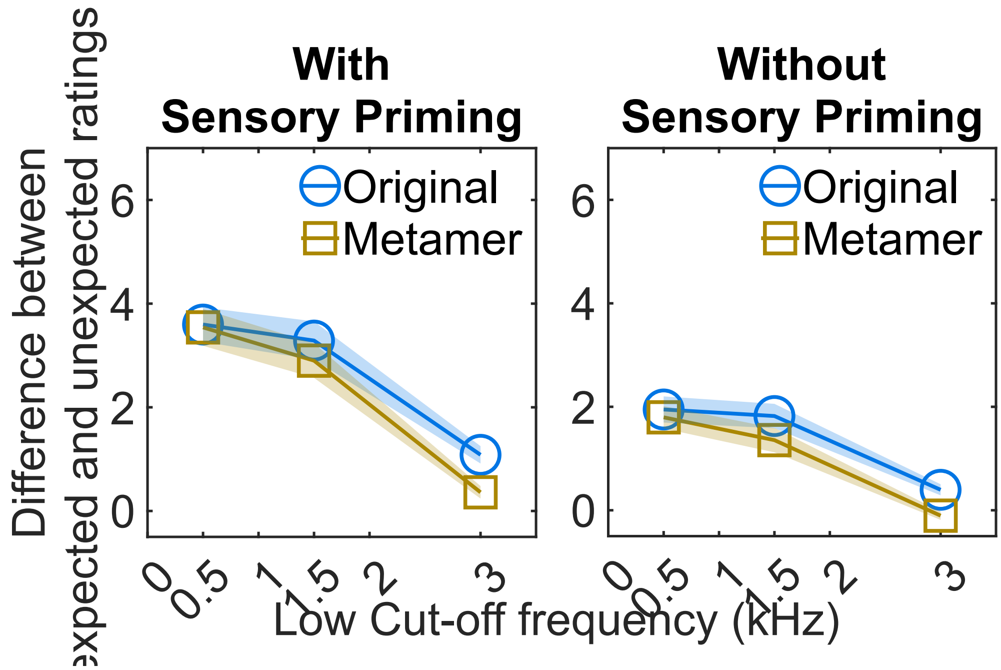


Supplementary Figure 4: Behavioral results from experiment 4, plotted as the difference between expected and unexpected ratings for each condition. Shaded regions ±1 standard error of the mean.

**Model comparison statistics**

We compared model responses using root-mean-square error between model predictions and observed behavioral responses. For Experiments 1-2 in each condition, we computed RMSe using values as originally measured for behavioral thresholds, in units of log2(semitones). For Experiment 3, units are the sensitivity index d'. These values are listed below by experiment.

**Table 1 (Experiment 1 RMSe values):**

| | **F0 = 200 Hz** | | | |
|---|---|---|---|---|
| **Lowest rank:** | **4.5** | **7.5** | **10.5** | **13.5** |
| **Rate-place ORIG** | 0.9404 | 0.8925 | 0.7645 | 1.0158 |
| **Rate-place META** | 1.1015 | 1.1639 | 0.3939 | 0.6181 |
| **Timing ORIG** | 2.4673 | 2.0956 | 2.6937 | 1.1893 |
| **Timing META** | 2.4627 | 2.9505 | 2.3737 | 1.6918 |
| | **F0 = 1400 Hz** | | | |
| **Lowest rank:** | **4.5** | **7.5** | | |
| **Rate-place ORIG** | 2.544 | 4.8489 | | |
| **Rate-place META** | 1.9455 | 5.4568 | | |
| **Timing ORIG** | 2.118 | 4.1654 | | |
| **Timing META** | 0.7465 | 3.6247 | | |

**Table 2 (Experiment 2 RMSe values):**

| | | | | F0 = 112.5 Hz | | | |
|---|---|---|---|---|---|---|---|
| **Low cutoff (kHz)** | **0.5** | **0.7** | **1** | **1.5** | | | |
| **Rate-place ORIG** | 1.2687 | 1.3293 | 0.7495 | 0.2756 | | | |
| **Rate-place META** | 1.1741 | 0.9501 | 0.6538 | 0.2503 | | | |
| **Timing ORIG** | 6.1873 | 6.5243 | 6.8953 | 7.1706 | | | |
| **Timing META** | 3.3077 | 3.7695 | 3.5599 | 2.9145 | | | |
| | | | | **F0= 300 Hz** | | | |
| **Low cutoff (kHz)** | **0.5** | **0.7** | **1** | **1.5** | **2** | **3** | **4** |
| **Rate-place ORIG** | 1.3189 | 1.271 | 1.1215 | 1.7683 | 1.4412 | 0.8074 | 0.0496 |
| **Rate-place META** | 1.3715 | 1.3871 | 1.4331 | 2.1644 | 1.5662 | 1.2736 | 0.2806 |
| **Timing ORIG** | 2.7252 | 3.1398 | 3.3541 | 2.2462 | 2.4353 | 2.0634 | 0.7849 |
| **Timing META** | 2.3331 | 2.8551 | 2.8516 | 2.509 | 2.6338 | 2.0618 | 0.9865 |
| | | | | **F0 = 900 Hz** | | | |
| **Low cutoff (kHz)** | | | | **1.5** | **2** | **3** | **4** |
| **Rate-place ORIG** | | | | 1.2652 | 1.1069 | 1.9477 | 1.1429 |
| **Rate-place META** | | | | 1.2292 | 1.2542 | 1.4471 | 1.2659 |
| **Timing ORIG** | | | | 1.649 | 2.0812 | 1.3755 | 0.7449 |
| **Timing META** | | | | 1.1984 | 1.9005 | 1.3571 | 1.2357 |

**Table 3 (Experiment 3 RMSe values):**

| | | | | F0 = 112.5 Hz | | | |
|---|---|---|---|---|---|---|---|
| **Low cutoff (kHz)** | **0.5** | **0.7** | **1** | **1.5** | | | |
| **Rate-place ORIG** | 1.1846 | 0.9598 | 0.9437 | 0.9146 | | | |
| **Rate-place META** | 1.0591 | 0.9472 | 0.9362 | 0.8634 | | | |
| **Timing ORIG** | 3.4119 | 3.4586 | 3.1448 | 2.6474 | | | |
| **Timing META** | 2.6311 | 1.8683 | 1.1947 | 0.9234 | | | |
| | | | | **F0= 300 Hz** | | | |
| **Low cutoff (kHz)** | **0.5** | **0.7** | **1** | **1.5** | **2** | **3** | **4** |
| **Rate-place ORIG** | 1.2729 | 0.9701 | 0.8388 | 1.1678 | 1.1259 | 1.0102 | 0.9473 |
| **Rate-place META** | 1.4241 | 1.176 | 1.0492 | 1.0695 | 0.9311 | 0.6247 | 0.7935 |
| **Timing ORIG** | 2.6507 | 2.7348 | 2.4562 | 2.0671 | 1.4322 | 1.8702 | 1.6805 |
| **Timing META** | 2.4278 | 2.4202 | 2.4545 | 1.9151 | 1.4518 | 0.8587 | 0.9788 |
| | | | | **F0 = 900 Hz** | | | |
| **Low cutoff (kHz)** | | | | **1.5** | **2** | **3** | **4** |
| **Rate-place ORIG** | | | | 1.2516 | 1.1061 | 1.1308 | 1.5058 |
| **Rate-place META** | | | | 1.3153 | 1.4883 | 1.011 | 1.6507 |
| **Timing ORIG** | | | | 1.1289 | 0.6736 | 0.9244 | 0.9483 |
| **Timing META** | | | | 1.0605 | 0.9142 | 1.189 | 0.553 |
